# An unexpectedly effective Monte Carlo technique for the RNA inverse folding problem

**DOI:** 10.1101/345587

**Authors:** Fernando Portela

## Abstract

Solving the RNA inverse folding problem, also known as the RNA design problem, is critical to advance several scientific fields like bioengineering, yet existing approaches have had limited success. The problem has several features that resist traditional computational techniques, such as its exponential complexity and the chaotic behavior of its cost function. Although some state-of-the-art AI approaches have reported promising results, all existing computational methods substantially underperform expert human designers. I combine a different technique, Nested Monte Carlo Search (NMCS), with domain-specific knowledge to create an algorithm that outperforms all prior published methods by wide margins and solves 95 of the 100 puzzles listed in a recently proposed RNA solving difficulty benchmark.

## Introduction

The RNA inverse folding problem is crucial in numerous scientific fields such as pharmaceutical research, synthetic biology and RNA nanostructures. Even though the question of the computational complexity of the RNA design problem is not categorically settled, recent evidence suggests it is NP-complete (Bonnet, Rzążewski & Sikora, 2017). Moreover, target-specific structural features like symmetries and short helices can heavily compound the difficulty of solving a particular RNA design problem (Anderson-Lee et al., 2016).

It is therefore unsurprising that existing RNA design software packages explore search spaces by way of trial and error. Examples of classic cost function minimization approaches include packages like: RNAinverse (Hofacker, 2003), performing adaptive random walk; RNA-SSD (Andronescu et al., 2004), using hierarchical structure decomposition; INFO-RNA (Busch & Backofen, 2006), probabilistic sampling of sequences; NUPACK (Zadeh et al., 2011), ensemble defect minimization; and MODENA (Taneda, 2015), a genetic algorithm. However, none of these packages come close to matching the performance of talented human RNA designers in the Eterna100 benchmark (Anderson-Lee et al., 2016): 54/100 for the best machine to 100/100 for the most talented human experts. Recent efforts using more sophisticated AI techniques include software packages like SentRNA (Shi, Das & Pande, 2018) (Eterna100 score 80/100) applying Deep Learning techniques incorporating a prior of human design strategies, and MCTS-RNA (Yang et al., 2017) (Eterna100 score 72/100) implementing a Monte Carlo Tree Search (MCTS) process largely inspired by computational treatments of the game of Go (Gelly & Silver, 2008) that were fashionable until the advent of DeepMind’s alphaGo (Silver et al., 2017).

The particular form of MCTS implemented in MCTS-RNA, called Upper Confidence Bounds applied to Trees (UCT) (Kocsis & Szepesvári, 2006), is known to be well suited for finding near-optimal solutions in huge solution spaces, themselves embedded in typically gigantic search spaces. Challenging RNA design problems often lack substantially large solution spaces though, especially when considered in relation to the sizes of their respective search spaces. The rarity of solutions within large subtrees effectively creates trap states (Ramanujan, Sabharwal & Selman, 2011) and causes the UCT search to ignore these subtrees for long periods of time (Mehat & Cazenave, 2010), making UCT an ineffective approach in such contexts. However, the field of General Game Playing (GGP) has produced more than one Monte Carlo algorithm. In particular, the simpler and well-studied Nested Monte Carlo Search (Cazenave, 2009) algorithm has been shown to be superior to UCT in many single player games (Mehat & Cazenave, 2010) and has never been tried in the context of RNA inverse folding.

Meanwhile, even though SentRNA still underperforms human solvers in the Eterna100 benchmark, its incorporation of human design strategies—in the form of a dataset of experts’ solutions to numerous RNA puzzles––shows potential. However, the solution to a given puzzle says nothing about *how* a human expert walked the tortuous path to the successful outcome. One could conjecture that the process itself, not just the final product, probably holds consequential and valuable information.

In addition, a quick survey of the newly collected move histories data on the Eterna game platform (Lee et al., 2014) had convinced me that several behavioral patterns in players’ solving styles could be encoded as algorithms. Combining all these observations, I hypothesized that implementing a Nested Monte Carlo Search (NMCS) based RNA inverse folding agent enhanced by heuristics in both the sampling and the explorative phases could lead to a best-of-class ability to solve RNA design problems that are intractable to current computational methods.

## Methods

The NEsted MOnte Carlo RNA puzzle solver (NEMO) is implemented as a short single C++ file. The linked RNA folding engine is the ViennaRNA package (Lorenz et al., 2011) in its version 2.1.9. A general overview of NMCS is presented in Fig. 1. NEMO’s simple global algorithmic layout is depicted in Fig. 2.

**Figure 1.**
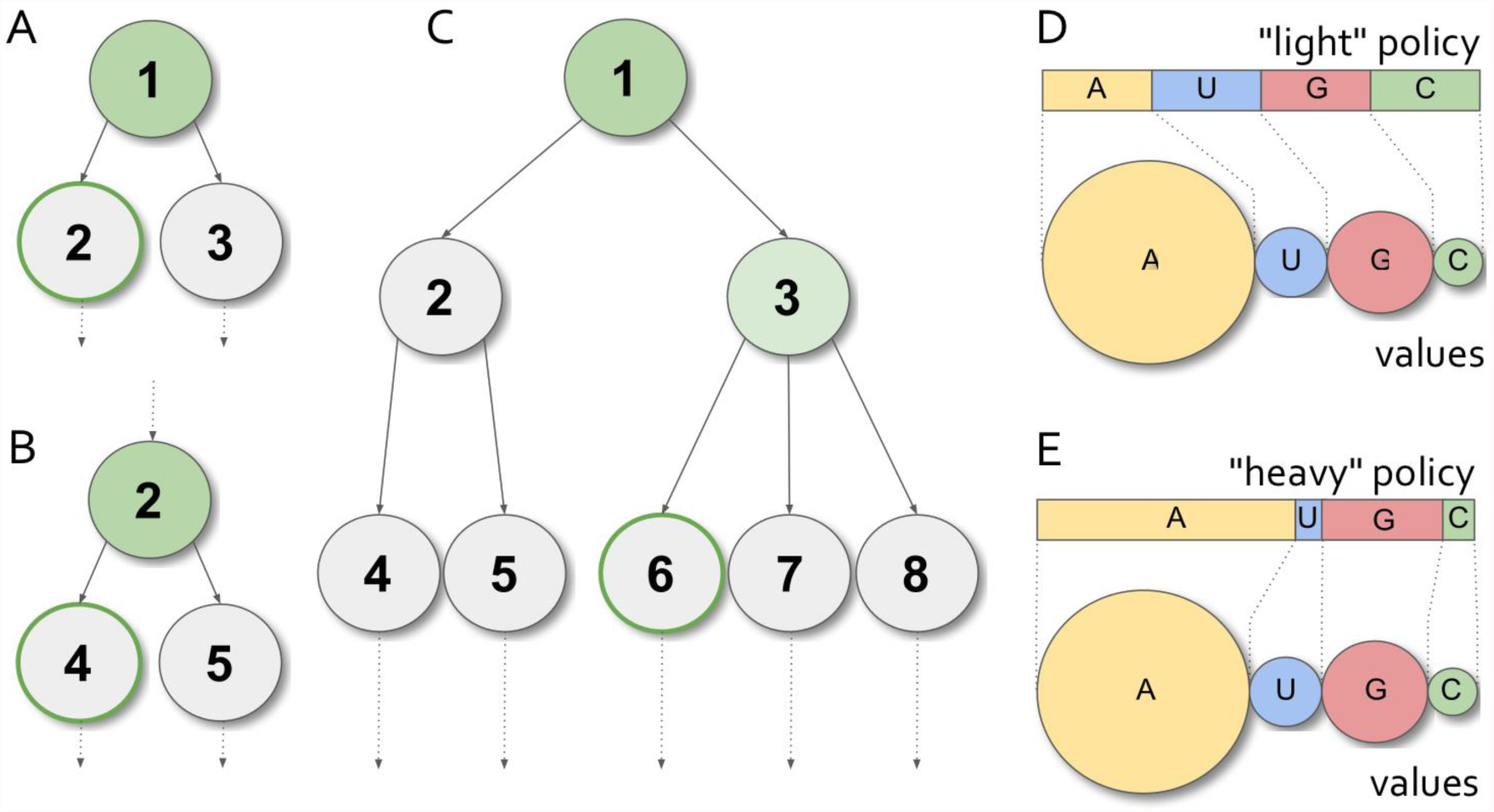
NMCS levels and playout policies. (**A**) Level-1 NMCS procedure: in an hypothetical game, state 1 has two legal moves, leading to states 2 and 3. The game is played out randomly with both options, the best scoring random game (here with state 2) is selected, and state 2 becomes the new root state. (**B**) The same procedure is applied until the game is finished. (**C**) The level-2NMCS procedure is similar and tests all grand-children nodes rather than the direct children ones. States 4 to 8 are all played out randomly. Here, state 3 becomes the root state for the next iteration. (**D**) Playout policies are applied to choices in the selection of a move in a Monte Carlo playout. Equiprobability is the simplest form of probability distribution. (**E**) The quality of the sampling can be strongly influenced by “heavy” playouts: using heuristics (which cost CPU resources), the software makes an educated guess as to which option is more valuable. The best playout policies are those that guess the correct order of preference in the available moves. In this example, the “heavy” playout policy makes a decent guess, even though it produces A>G>C>U when A>G>U>C would have been preferable.

**Figure 2.**
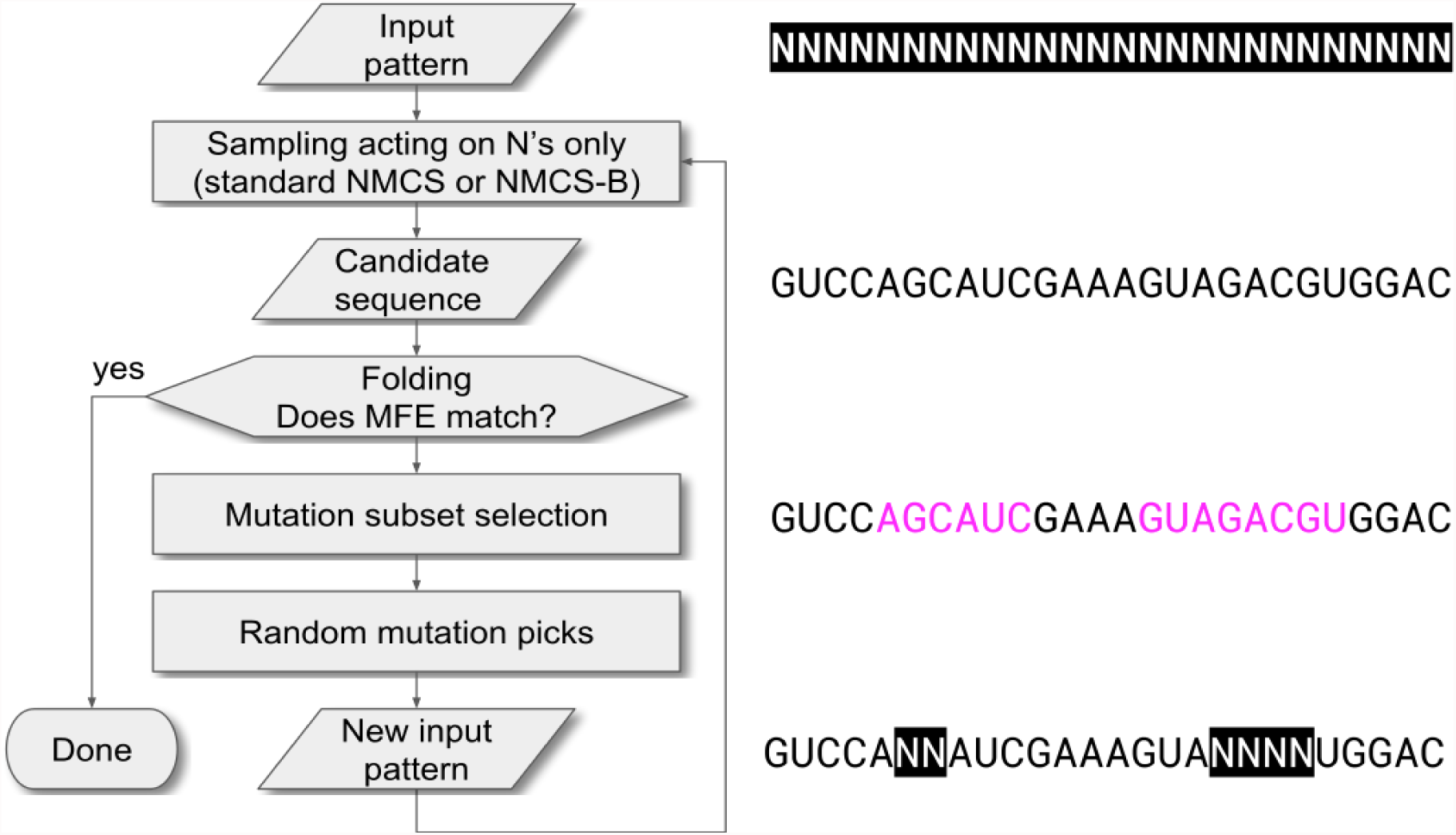
Schematic of NEMO’s algorithm with an example of the evolution of the internal state over the first iteration.

### Sampling phase heuristics

The heuristics and strategies in the sampling phase—the playout policy in General Game Playing theoretic parlance—are coded using domain knowledge acquired by personal experience. Parameters affecting probabilities and distributions were chosen ad hoc, without performing computational optimizations, whether gradient descent or otherwise. Its initial step consists in filling up base pairs first, and only then the unpaired positions in the target structure. Following this order allows NEMO to properly handle specific sub-goals like preventing unwanted base pairings in 0-N bulges—a technique known as “blocking” among Eterna players—and using thermodynamically favorable mismatches—also known as “boosting”—in multi-way junction loops. (Note: In this paper, the term “mismatch” is meant to include all form of potential non-canonical interactions at the end of helices, e.g. both terminal mismatches and dangling ends)

Roughly following the proportions found in known naturally occurring RNA structures (Lemieux & Major, 2002), NEMO fills base pairs with a 60% GC, 33% AU and 7% GU probability distribution, with a few exceptions for closing pairs of adjacent helices in junctions and the closing/enclosing pairs of triloops. Unpaired bases are divided into two categories depending on whether they participate in mismatch interactions or not. Since non-mismatched bases are thermodynamically neutral in the Turner model (Turner & Mathews, 2010), their nature should not be a concern, but in practice common puzzle-solving wisdom suggests to make these domains A-rich; for these bases, NEMO uses a 93% A, 1% U, 5% G and 1% C probability distribution. Mismatched bases however do affect the Gibbs free energy contributions of loops and are therefore highly relevant for finding solutions. In such cases, NEMO uses heuristics (described in Fig. 3) derived from <u>Eterna game playing experience</u>.

**Figure 3.**
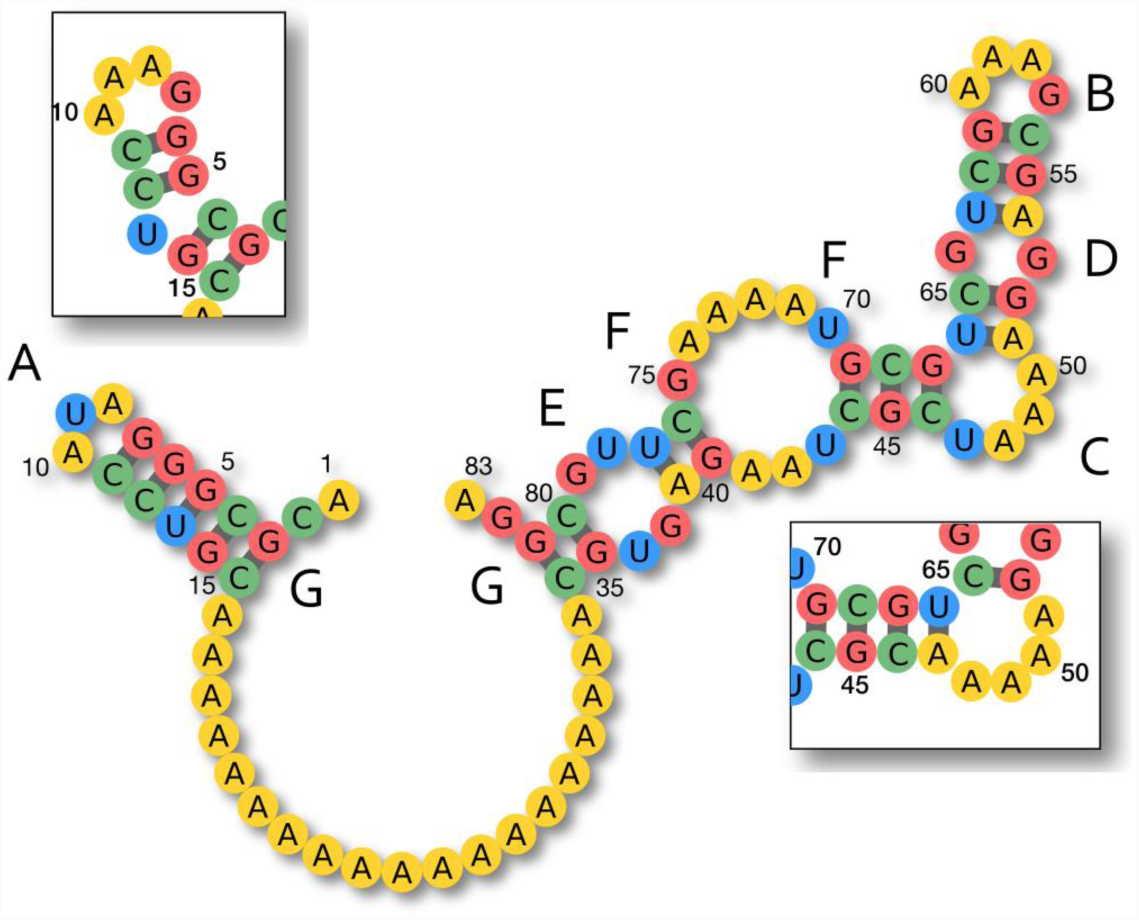
Playout policy heuristics for unpaired mismatched bases. (**A**) “blocking” trick occasionally required or simply helpful in some triloops, the inset demonstrates the “sliding” that happens with the U9A mutation (**B**) standard G/A mismatch in apical loops (**C**) “blocking” applied to a bulge, the inset demonstrates the unwanted pairing occurring with the U47A mutation (**D**) standard G/G mismatch in symmetric 1-1 internal loops (**E**) standard UG/UG combo-mismatch in symmetric 2-2 internal loops (**F**) some typical favorable mismatches (“boosts”) in internal loops, here A/G and U/U (**G**) favorable mismatches for external loops and junctions, C/A for GC closing pairs, A/G for CG closing pairs.

### Cost function

The scoring of the samples is a composite function of:

• the base pair distance (BPD) between the Minimum Free Energy (MFE) structure of the sample sequence as calculated by the folding engine and the target structure, expressed as 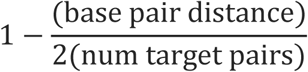

• and the ΔΔ*G* between the MFE of the sample sequence, and its predicted Gibbs free energy in the target conformation, expressed as 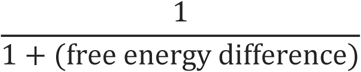

### NMCS variants

Two slightly different versions of the NMCS algorithm were implemented: the standard version as found in the GGP literature, and a modified one—that I labeled NMCS-B, standing for “Nested Monte Carlo Search with Best playout policy”—where I introduced an internal maximization mechanism that retains the best scoring playouts throughout the recursion (the difference between them is shown in Fig. 4). Both were executed at the standard level 1 of recursion. Using NMCS recursion at levels 2 or higher would significantly increase computational cost (by up to 30-fold) and was not tested.

**Figure 4.**
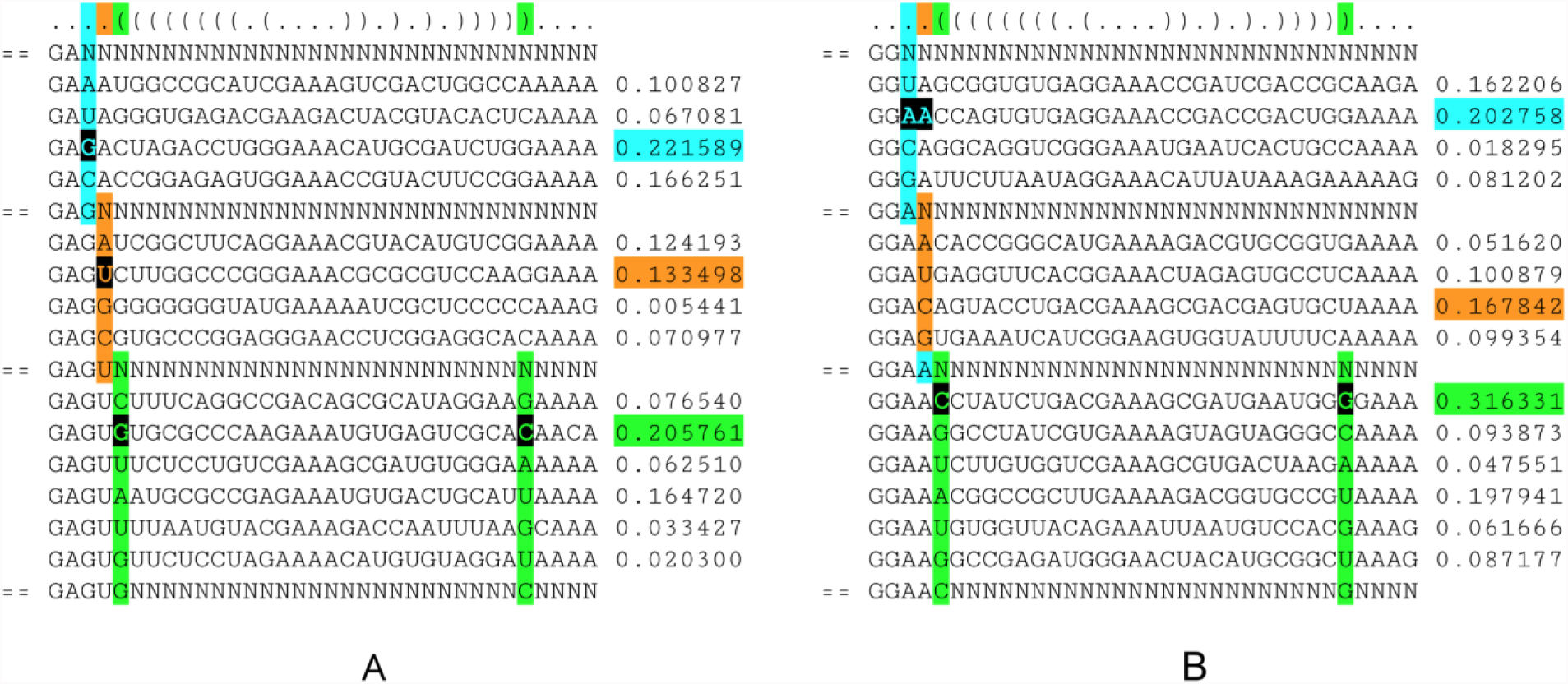
Algorithmic difference between standard NMCS and NMCS-B. Top row, the target structure in dot-bracket notation. In both cases, the algorithm processes positions in order: cyan, orange, green, etc. (A) “vanilla” Nested Monte Carlo Search (NMCS) considers all possible choices at each position by doing one Monte Carlo sampling, and always picks the best of these outcomes. (B) NMCS-B differs in that it retains the best playout so far and compares it to the samples generated at each recursion step. Here NMCS-B ignores the option C at the orange position, because a better cyan sample playout is known. At the green step, the sample playout sporting the CG pair becomes the best sample known so far and might influence the next steps.

### Selection heuristics for iterations

After the evaluation of a candidate sequence, and provided it was a failure, NEMO identifies the subset of the sequence for which mutations should be considered in preparation for the next iteration by first collecting the indices of all the bases that didn’t fold as expected, and then expanding this set with other potentially relevant indices: first adding all mismatch partners of the already collected misfolded positions (as pictured in Fig. 5) and then including closing pairs neighboring pairs that are misfolding by “opening up” (as described in Fig. 6).

**Figure 5.**
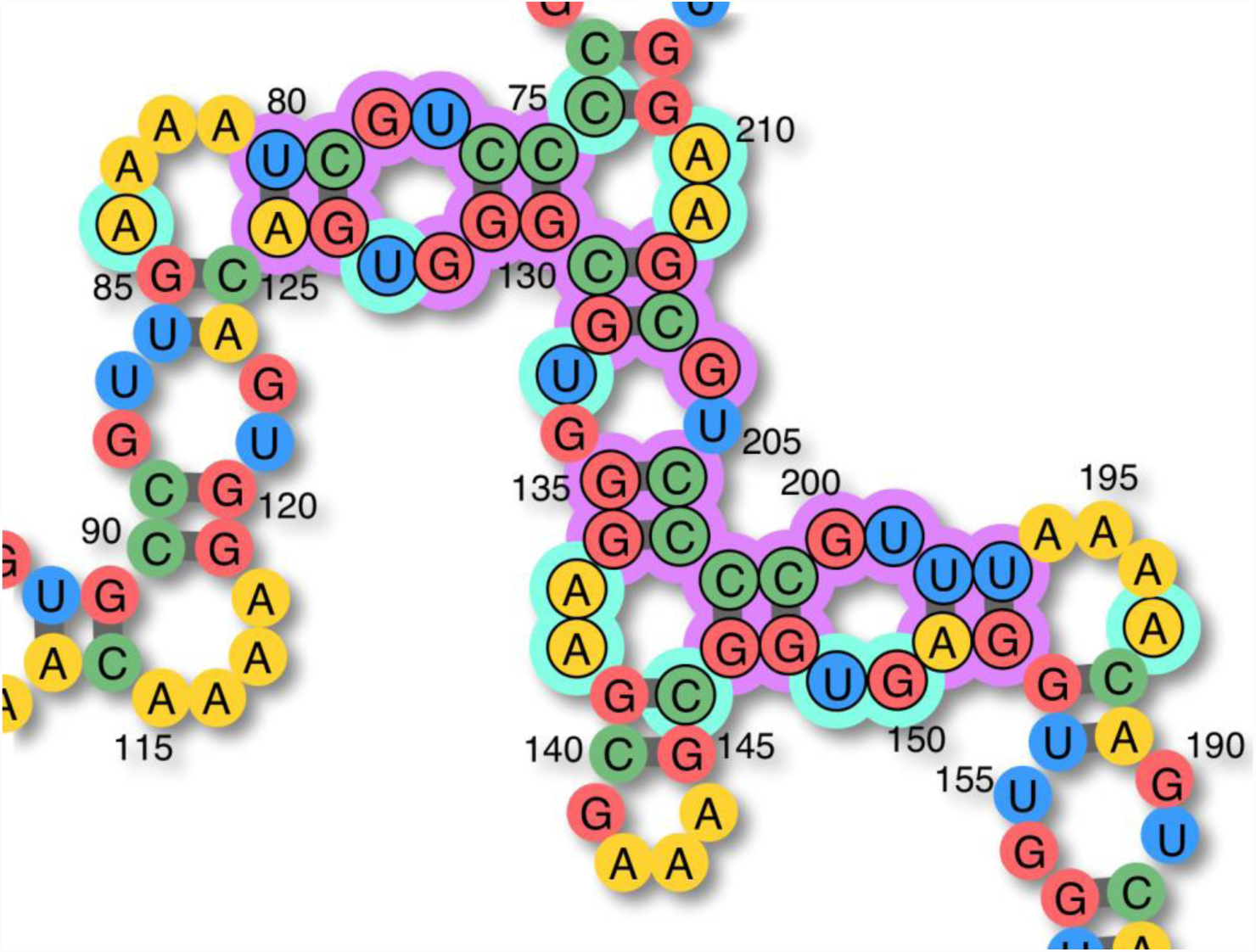
Mutation candidates subset selection. This program phase begins with including all misfolded bases (purple marks). Additionally, their mismatch partners (cyan marks) are included as well. Note: in this example (which has a base pair distance of 24 w.r.t. the target structure), the A84G and A210C mutations both stabilize the puzzle completely. In other words two solutions exist only 1 one-point mutation away from this particular sequence.

**Figure 6.**
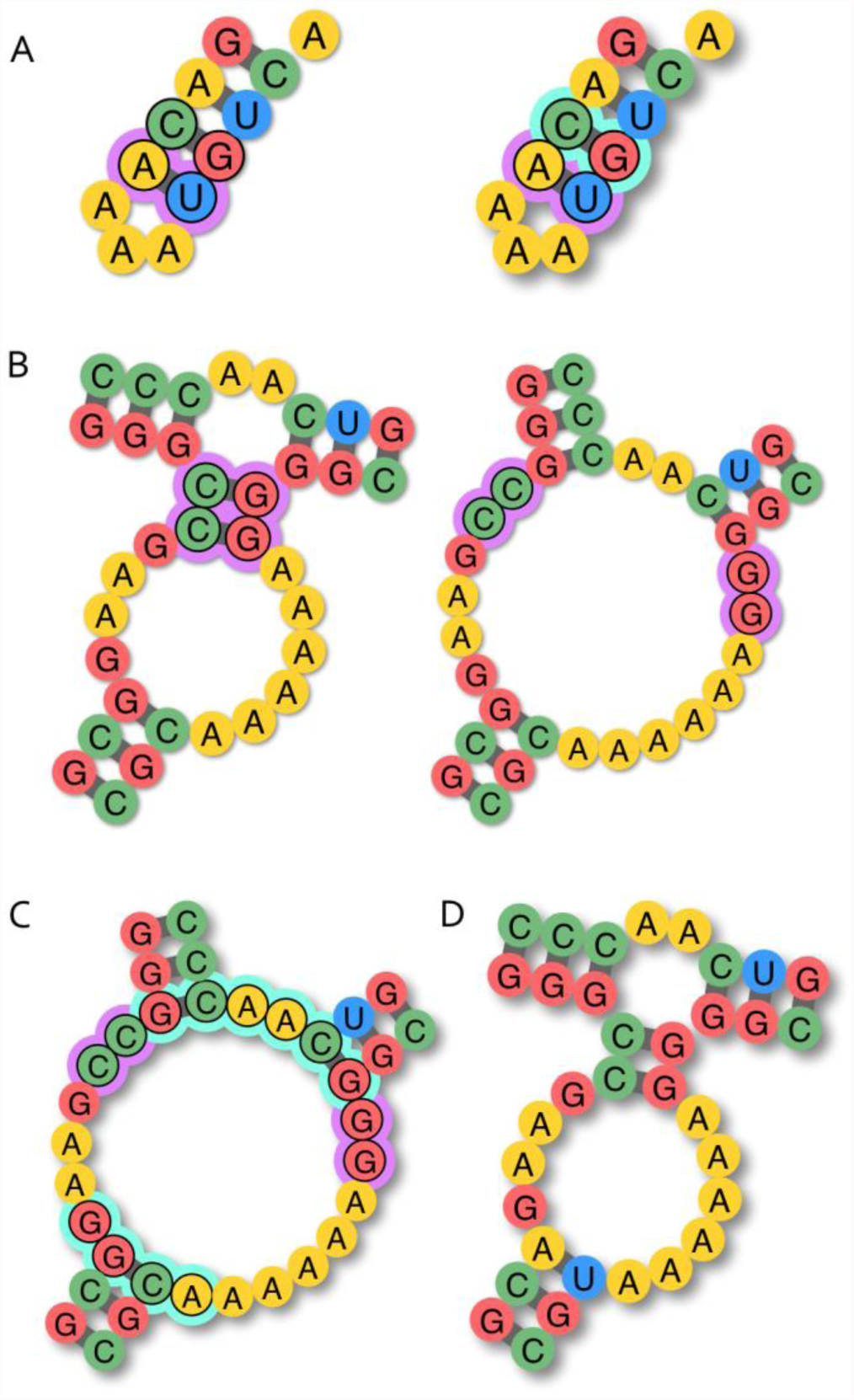
Heuristic rule of including closing pairs around pairs misfolding by “opening up”. (**A**) An AU pair closing a triloop cannot hold if the enclosing pair is GC/CG. Mutating the closing pair to CG/GC would likely solve the issue, but at this stage, the goal is to collect all mutations that could. Changing the enclosing pair to UA could solve the local problem as well, therefore the cyan marked pair should be included in the subset. (**B**) A slightly more complex case with a short stem linking two loops. (**C**) The pairs closing the large junction in the misfolded structure, and their associated mismatches (cyan marks) should all be added to the mutation candidates list. (**D**) In this particular case, it turns out that no mutations of the purple-marked pairs and no alternate boosting of the surrounding loops can help the short helix to hold in place. The only mutation that works here, is to change the bottom-left closing pair to AU/UA.

### Testing

In order to test the algorithm and measure its fitness, I repeatedly ran the NEMO tool against the Eterna100 benchmark (Anderson-Lee et al., 2016). Performance and success rates of various builds were measured over 30 single-shot batch runs executed on Stanford University’s BioX^3^ and Sherlock clusters. Each process had a default limit of 2500 iterations, corresponding to a maximum of approximately 90 minutes on a single Intel^®^ Core^TM^ i7 3.1 GHz processor for a 400 nucleotides long design problem. Separately, MCTS-RNA and NEMO were both tested in the precise conditions used in (Anderson-Lee et al., 2016): up to 5 attempts spanning a maximum of 24 hours.

## Results

### Self comparisons

Average iteration counts and success rates for the comparison between standard NMCS, NMCS-B and weakened versions of NMCS-B are presented in Fig. 7. The raw data are provided in the Supporting Documents.

**Figure 7.**
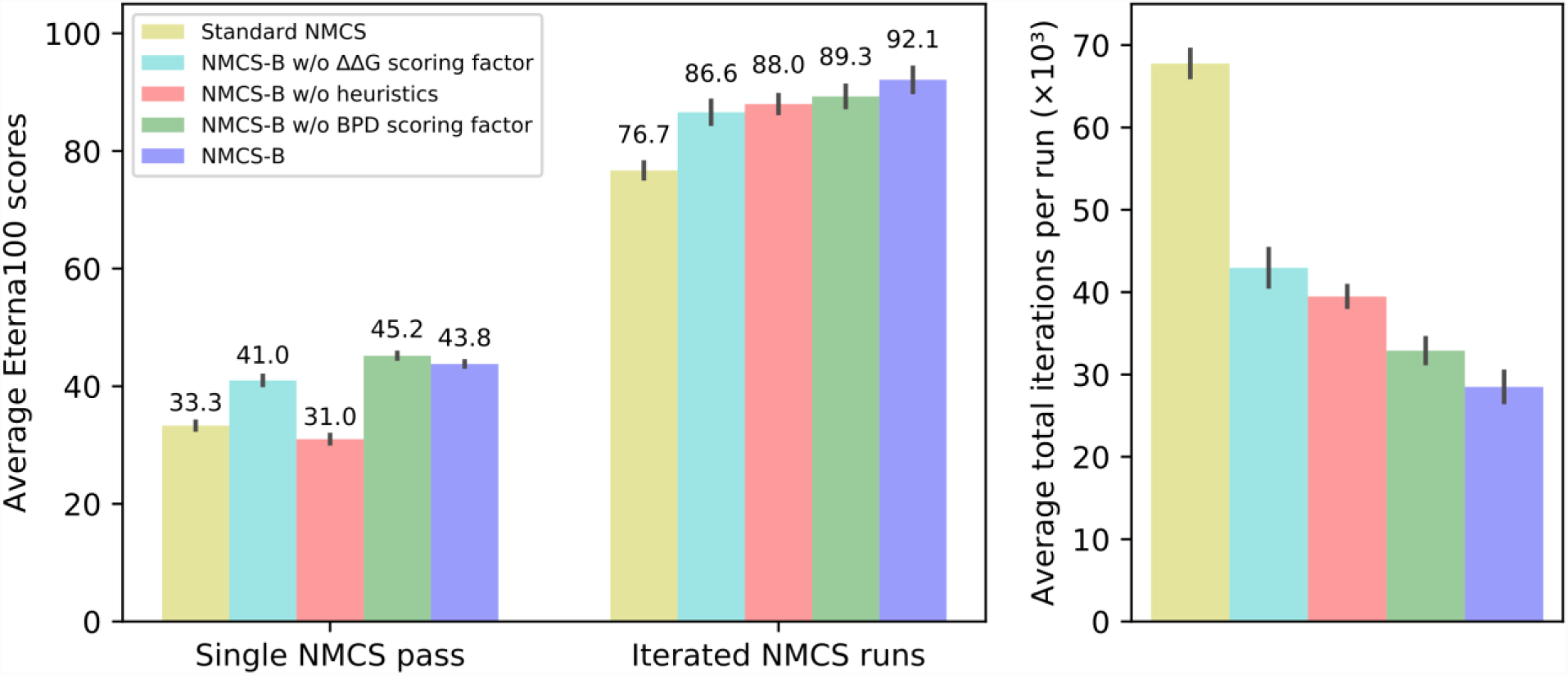
NEMO performance tests. Average scores and iteration counts over 30 single attempts runs against the Eterna100 benchmark.

NMCS-B demonstrates a clear superiority over the standard version of NMCS, both by solving about 15 more puzzles on average (92.1/100 to 76.7/100), and by converging to solutions more than twice as fast. As expected, removing algorithmic elements from NEMO causes the performance to worsen: decreasing by about 3 to 6 points in puzzle-solving power, and significantly slowing down convergence by between 15% and 50%. The average success rate of the first NMCS-B pass alone (e.g. solution found with no iterations) is only 43.8/100 though.

### Comparisons with other engines

When coupled with the standard form of NMCS, the performance of NEMO (77/100) compares to that of the best solving engines previously tested on the same benchmark, the top contender so far being SentRNA with 80/100. But when using the NMCS-B variant, NEMO surpasses them all by a comfortable margin, solving 95 puzzles out of 100 (Fig. 8), and clearly outclasses the other bandit-based method (UCT) implemented in MCTS-RNA which scored 72/100.

**Figure 8.**
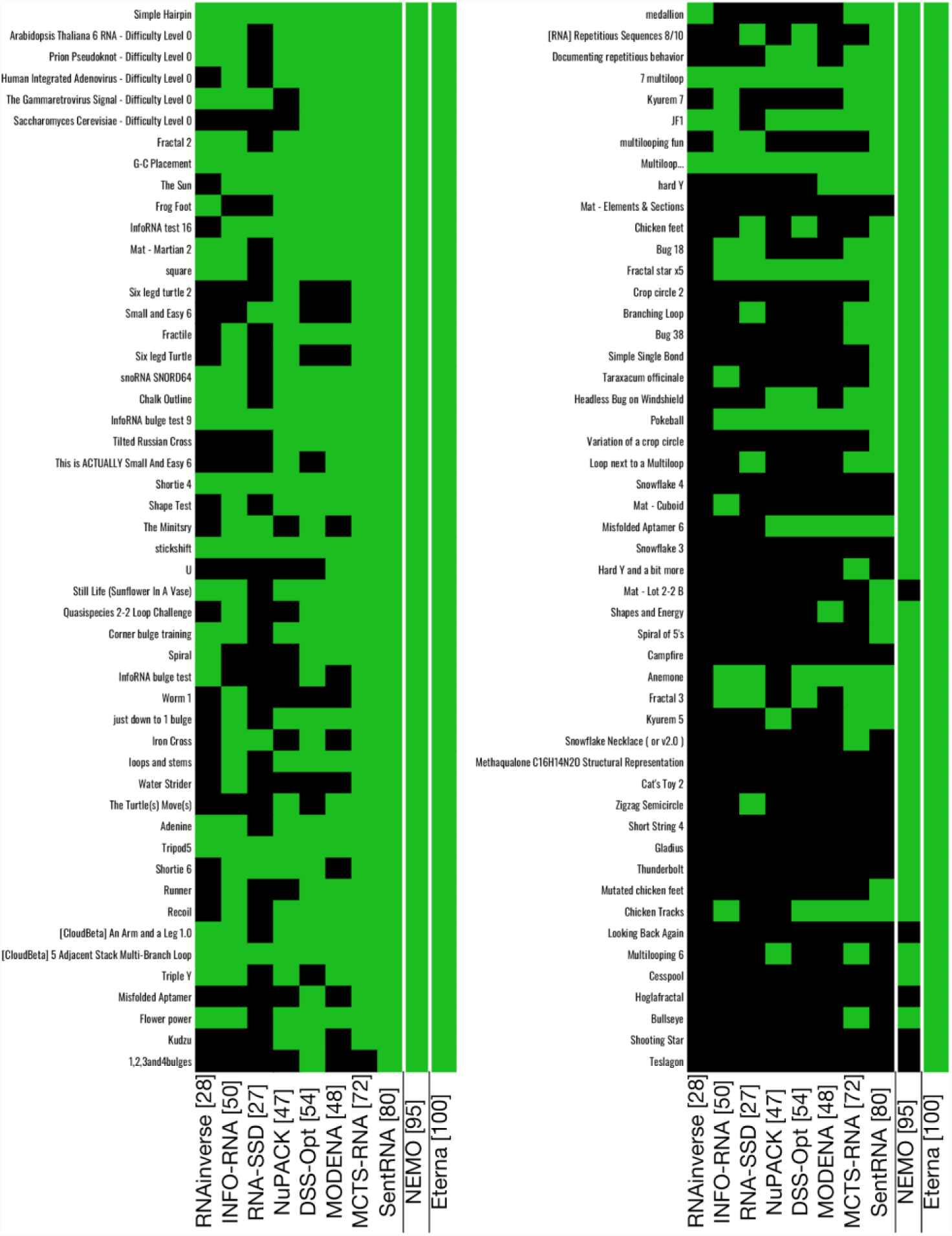
Comparative Eterna100 benchmark results. Black squares are failures, green ones are successes. RNAinverse, INFO-RNA, RNA-SSD, NuPACK, DSS-Opt, MODENA et Eterna players results data were taken from Anderson-Lee et al. 2016, SentRNA data taken from Shi et al. 2018. MCTS-RNA was configured to ignore GC content requirements. NEMO was run with NMCS-B active.

## Discussion

### Unexpected effectiveness

The results point to an overall excellent fitness of the NMCS algorithm when applied to the RNA inverse folding problem. However, the reasons for NEMO’s strong performance against the Eterna100 benchmark are not entirely clear.

The argument that it would be exclusively linked to the quality of the domain knowledge integrated into the tool is at odds with the fact that NEMO only has a limited set of helpful heuristics, lacking for instance special code for dealing with many other known RNA structural complexities like “zigzags” (example depicted in Fig. 2f of (Anderson-Lee et al., 2016)). Furthermore, a direct test by depriving NEMO of heuristics reduces its performance only slightly.

Also, even though the scoring function in NEMO appears to be novel (Gibbs free energies have been used as selection criterion (Hampson, Sav & Tsang, 2016) but in absolute rather than relative terms) compared with those implemented in other RNA design software packages, its output smoothness over the search spaces remains insufficient to possibly explain any substantial sensitivity improvement in the exploration process, which is supported by the data showing a definite but only modest improvement from using ΔΔ*G*.

Finally, the effectiveness of NMCS versus UCT is indeed known in the General Game Playing field to be game-dependent (Mehat & Cazenave, 2010). Though the poor score of NEMO’s first NMCS phase taken alone in the Eterna100 benchmark test also contradicts the hypothesis that the RNA inverse folding game could simply be an excellent fit for the NMCS algorithm.

### Imitating human RNA designers

One possible explanation for NEMO’s performance is that it partially imitates the solving style of some of the successful human players on the Eterna game platform. In broad terms, the two main classes of puzzle-solving styles delineate a global-local dichotomy perceptible in recorded move histories. Thanks to the fact that the ΔΔ*G* can only be measured when the target structure is entirely filled with valid base pairs, the paths lengths in Fig. S1B allow deriving behavioral information. For instance, the Eterna players who produced the solutions 2, 5, 7 and 9 had filled their canvas early with valid pairs everywhere, while those who generated solution 1, 3 and 4 didn’t solve the puzzle any faster than the others, but rather favored an incremental method by dividing the problem into smaller ones and by stabilizing first each subdomain before tackling the next one. For that reason, their ΔΔ*G* “tracks” are much shorter than others. Since NEMO uses the ΔΔ*G* in its scoring function, it needs to mimic the “globalist” approach. As for mutating bases or reorienting pairs within or near the misfolded domains, this behavior is common to all Eterna players, and NEMO roughly imitates it in the subset-defining and random-picking phases done in preparation of the next iteration

As the data show, the NMCS procedure alone is often insufficient to provide an immediate solution to RNA puzzles, which parallels the fact that even expert Eterna players are unlikely to solve a hard puzzle in a single shot. For instance, only 45 players out of the 250,000 registered on Eterna (as of 2018) have solved the <u>“Snowflake 4” puzzle</u> in the Eterna100 benchmark. The fastest solver on record still required 11 minutes and 345 mutations to complete the challenge, almost 200 more operations than is needed to produce a minimally valid sequence for this puzzle. However, the purpose of the NMCS phase in NEMO is not to solve hard puzzles instantaneously, but to provide reasonable candidate sequences for the overlying explorative process. Qualitatively superior samples tend to benefit any Monte Carlo approach, provided their computational costs stay reasonable and on the condition that the introduced biases leave the relative weights of the visited nodes and subtrees mostly unaffected (James, Konidaris & Rosman, 2017). I encoded into NEMO parts of the domain knowledge (Fig. 3) I acquired by practice and by reading numerous <u>guides authored by fellow Eterna players</u>, and as evidenced by the data presented here, it enhances its NMCS playout policy.

The decision to combine base pair distances and ΔΔ*G* in the scoring function was both a reflection of my personal puzzle-solving style and the result of analyses on player-submitted solutions and their move histories. Figure S1 conveys that the base pair distance measurement, which is the measurement of choice for the vast majority of RNA inverse folding packages, is a turbulent variable that stays chaotic arbitrarily close to the end goal (an example of which is depicted in Fig. 5). In contrast, no matter the players’ puzzle-solving style, global or incremental, the ΔΔ*G* measurement seems much more reliable as an indicator for approaching a solution. The lower performance of the ΔΔ*G* scoring alone (without base pair distance) was predictable: RNA strands can reach multiple conformations, and without a solid structural indicator, an RNA solving engine can waste iterations chasing after a flawed construct that keeps tending to fold into alternate conformations. NEMO currently has no routines to perform local free energy optimizations, but incorporating the ΔΔ*G* in the global scoring function as a cofactor of the base pair distance presumably helps guide the search in the right direction, and the data support its appreciably positive effect.

During adaptive random walks or tree explorations in RNA puzzle-solving, simple common sense prescribes to keep the parts that fold correctly and only mutate the bases and pairs belonging to domains that do not. RNA inverse folding engines follow this guideline when their main measuring stick is the base pair distance, just as human experts generally do: for instance in the move histories of solutions to the Eterna100 benchmark puzzle titled <u>“Methaqualone C_16_H_14_N_2_O Structural Representation”</u>, expert players mutated misfolded bases 82.8% of the time on average (data provided in Supporting Documents). In that particular phase of the solving process, NEMO’s locality-based heuristics represent a crude but effective approximation of this human behavior.

However, the current implementation of NEMO lacks a mechanism for imitating a prominent behavior of successful RNA puzzle solvers: backtracking. Data collected on the same previously mentioned Eterna100 puzzle also indicate an average backtracking rate of 22.3%. Presumably, Eterna game players would oftentimes find themselves at a stage of the puzzle-solving process that they regard as “close to solving”. They would then carefully explore various branches of possible mutations, and undo their unsuccessful changes to come back to the previously found satisfactory state if the test was inconclusive. A similar behavior could be implemented in NEMO by replacing the iterated random walk by a judiciously crafted form of tree search.

### Generality of approach

Concrete applications of the RNA design problem usually require additional constraints like a specified GC content ratio, typically to precisely control melting temperatures for experiments using amplification by polymerase chain reactions (PCR) (Saiki et al., 1988). I intentionally ignored this specific goal in this work so as to better focus on the primary one: solve challenging RNA puzzles. The GC content control goal is trivial to achieve in a post-processing phase. Examples of such algorithms, which explore the neighborhood of a given solution and gradually change its GC/AU/GU pairs ratio, already exist in EternaScripts written by Eterna players (<u>“Jnicol`s - Remove the GCs v2” by mat747</u>).

In contrast, designing riboswitches is oftentimes a dissimilar endeavor. Structural constraints usually apply only to small domains within the design space, like binding sites for ligands or oligonucleotides, signaling domains, and gene expression initiation sequences. Structural freedom is granted for the rest of the construct. Should the need arise for a riboswitch with two (or more) precise target conformations, for instance for a nanostructural application, NEMO would have to be modified to properly handle “chain reactions”, i.e. the causal cascade of purines and pyrimidines pairing with different partners over multiple target structures. I already implemented such an algorithm in a previous work, a puzzle-solving bot (<u>Eterna profile of ViennaUCT)</u> competing on the Eterna game platform.

### Conclusion

Until recently, the potential of Monte Carlo techniques applied to the RNA design problem had remained mostly unexplored. The first implementation of the UCT algorithm in this context achieves a good performance against a collection of unyielding RNA puzzles that even human solvers struggle to complete. Though, the Nested Monte Carlo Search algorithm, enhanced by heuristics, outperforms both UCT and all other in silico RNA design approaches. Given the presented encouraging results, the Nested Monte Carlo Search, combined with a novel cost function formula and with a large extent of the accumulated knowledge of Eterna’s expert RNA designers, appears to be a promising technique worthy of deeper investigation by RNA design package creators.

## Data and materials availability

The source code of NEMO is available at https://simtk.org/projects/nemo. Raw tests results and moveset analysis are available at https://doi.org/10.6084/m9.figshare.6358625.

## Acknowledgments

The author would like to thank Rhiju Das, Benjamin Keep and Michelle Wu for their help with the manuscript, and Stanford University for granting him access to the BioX^3^ and Sherlock clusters.

## Supplementary Figures

**Figure S1.**
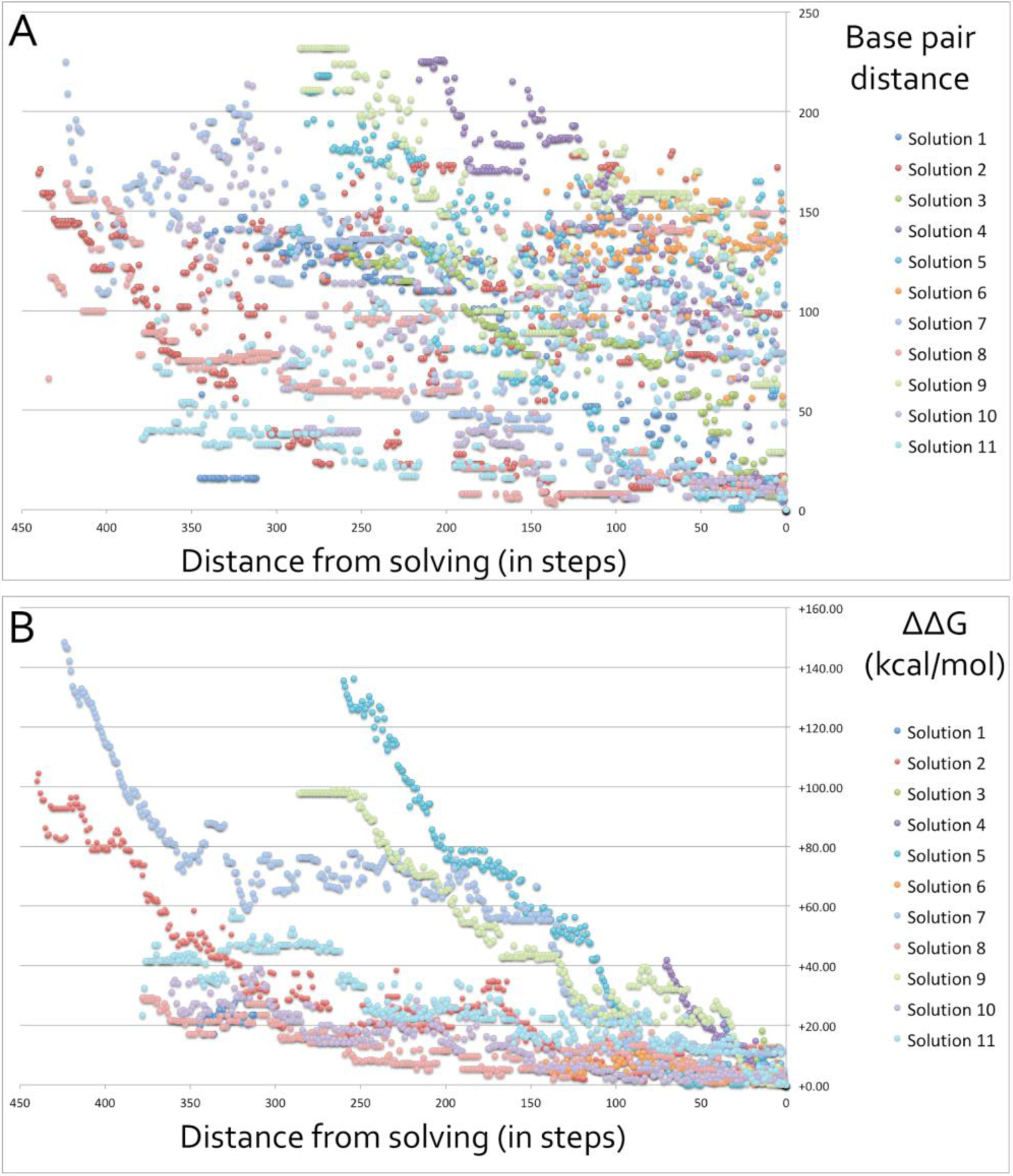
Base pair distances and free energy differences as functions of distance in mutation steps in human solving. Both graphs relate to solutions provided by human experts for the <u>“Snowflake 4” puzzle</u> of the Eterna100 benchmark. Evolution over time (in mutation steps) of (**A**) base pair distances, and of (**B**) Gibbs free energy differences (ΔΔ*G*) until a solution is found.

